# CapTrap-seq: Advancing zebrafish transcriptomic research through high-fidelity full-length RNA sequencing

**DOI:** 10.1101/2025.05.12.653332

**Authors:** Monika Kwiatkowska, Marta Blangiewicz, Tomasz Mądry, Silvia Carbonell-Sala, Roderic Guigó, Barbara Uszczynska-Ratajczak

## Abstract

Zebrafish is a valuable model organism thanks to its genetic and anatomical similarities to humans, offering a more relevant system for studying human biology and disease than *in vitro* or non-vertebrate models. However, its use in large-scale transcriptomic research is still limited. Most zebrafish studies focus on global expression profiling in specific developmental contexts, contributing little toward improving and expanding gene annotations. These gaps lead to inaccuracies in gene quantification and downstream functional analyses, ultimately reducing the effectiveness of zebrafish as a model system. Further challenges include ineffective ribodepletion methods and limited resources for validate transcript boundaries and splicing patterns. To overcome these challenges, we applied CapTrap-seq, a long-read sequencing method that combines Cap-trapping with oligo(dT) priming to identify 5’-capped, full-length transcripts from both developmental and adult tissue samples. To promote detection of longer RNA molecules, we introduced a size-selection step into CapTrap-seq protocol, which further improved detection of transcript ends without compromising its quantitative accuracy. Our results highlight the genome-agnostic nature of CapTrap-seq, enabling generation of accurate transcript models in non-mammalian systems at high-throughput. CapTrap-seq also facilitates functional annotation of zebrafish genes by uncovering novel, spliced transcript isoforms with potential biological significance.

## INTRODUCTION

Model organisms are non-human species that are invaluable for investigating the biological functions of genes and mimicking human disease symptoms through experimental or genetic manipulations that are not feasible in humans^1–3^. Mammalian models like mice, rats and primates have historically been favored for their complex biology and close evolutionary relationship to humans^4–7^. However, these models are expensive, less practical for large-scale genetic studies, and pose ethical concerns, particularly for research on early developmental stages. Zebrafish (*Danio rerio*), a small tropical fish native to South Asia, has emerged as a widely recognized alternative that complements the limitations of mammalian models^8–10^. Its advantages include low maintenance costs, ease of genetic manipulation, external development, transparency of embryos, and high molecular similarity to humans, with comparable major organs and tissues. As a result, zebrafish have become a powerful model for uncovering gene functions^11,12^ and studying molecular processes related to human biology^13–15^ and disease^8,9^. Despite these strengths, the use of zebrafish for large-scale gene discovery and transcriptomic studies remains largely underdeveloped. This limited application contrasts with their recognized value and restricts their contribution to systematic functional annotation efforts. Such efforts are crucial for uncovering the biological roles of protein-coding and non-coding genes and for understanding how their expression and splicing influence human health and disease^16–19^.

Due to economic constraints, zebrafish RNA sequencing (RNA-seq) have traditionally focused on specific biological contexts and primarily rely on standard short-read data^20,21^. Unlike long-read RNA-seq, short-read approaches face challenges in accurately reconstructing complex transcriptomes^22,23^ and detecting alternative isoforms^24,25^. Additionally, zebrafish RNA-seq experiments are typically performed at lower sequencing depths compared to human and mouse studies (50 vs 200-250 million reads)^26–28^. This limited coverage reduces transcriptome resolution and makes it difficult to reconstruct accurate transcript models at the single-molecule level, particularly for low-abundance genes or those with complex splicing^29^. As a result, zebrafish gene annotations remain incomplete, with many transcripts either partially annotated or entirely missing^30^.

A further limitation is the absence of large-scale systematic functional annotation programs for zebrafish, analogous to ENCODE or FANTOM for human, which restricts access to comprehensive genomic and epigenetic datasets needed to validate and interpret experimental findings. The recent release of DANIO-CODE data^31,32^ has improved the interpretability of zebrafish genome activity. However, this resource is largely focused on early development. While some zebrafish transcriptomic studies include adult samples, most remain focused on embryonic stages^20,33^. In contrast, human transcriptomic datasets emphasize adult tissues^26^, making zebrafish less useful for studies involving adult gene expression or tissue-specific regulation.

To address current limitations and advance the use of zebrafish in large-scale transcriptomic and functional studies, we applied our recently developed CapTrap-seq^34–36^ method combined with Oxford Nanopore Technologies (ONT) long-read sequencing to generate full-length transcripts from both developmental and adult tissue samples. When benchmarked against a template-switching protocol, CapTrap-seq demonstrated the ability to resolve annotation gaps and produce accurate transcript models for both protein-coding and non-coding genes beyond mammalian systems. Integrating a size-selection step further improved sensitivity for detecting complete transcript boundaries. A key strength of CapTrap-seq in zebrafish is its effective elimination of ribosomal RNA (rRNA), addressing a long-standing challenge due to limited rRNA depletion options for this species. In addition, CapTrap-seq has substantially expanded the limited catalog of alternatively spliced isoforms, providing a deeper view of transcriptome complexity and enabling more detailed functional annotation of zebrafish genes.

## RESULTS

### CapTrap-seq for full-length transcript identification in zebrafish

We applied our newly developed CapTrap-seq protocol^34,35^, which integrates Cap-trapping and polyA tailing with long-read cDNA sequencing (Figure 1A), to identify full-length transcript molecules from two early zebrafish developmental stages (2–4 cell, RIN=9.2 and 28 hpf, RIN=9.3) and two transcriptionally complex adult tissues (heart, RIN=8.9 and testis, RIN=9.2) (Figure 1B). To evaluate CapTrap-seq performance, we compared it with a standard library preparation method based on the Template-Switching Oligo (TSO) approach^37^ (Figure 1A). All samples were sequenced using the ONT platform, and the resulting reads were processed with an in-house LyRic pipeline^38^ to generate a non-redundant set of transcript models (TMs) per sample. In our previous study^34^, we successfully evaluated CapTrap-seq at the read level to minimize potential biases from downstream analysis. Here, we shift focus to transcript-level benchmarking, as transcript completeness is essential for accurate gene annotation, particularly for less extensively studied species such as zebrafish. We evaluated the completeness of LyRic-inferred TMs by analyzing the presence of unmapped poly(A) tails at their 3′ ends and their proximity to CAGE tags (N=1,585,796) from the DANIO-CODE project^31,32^.

**Figure 1.**
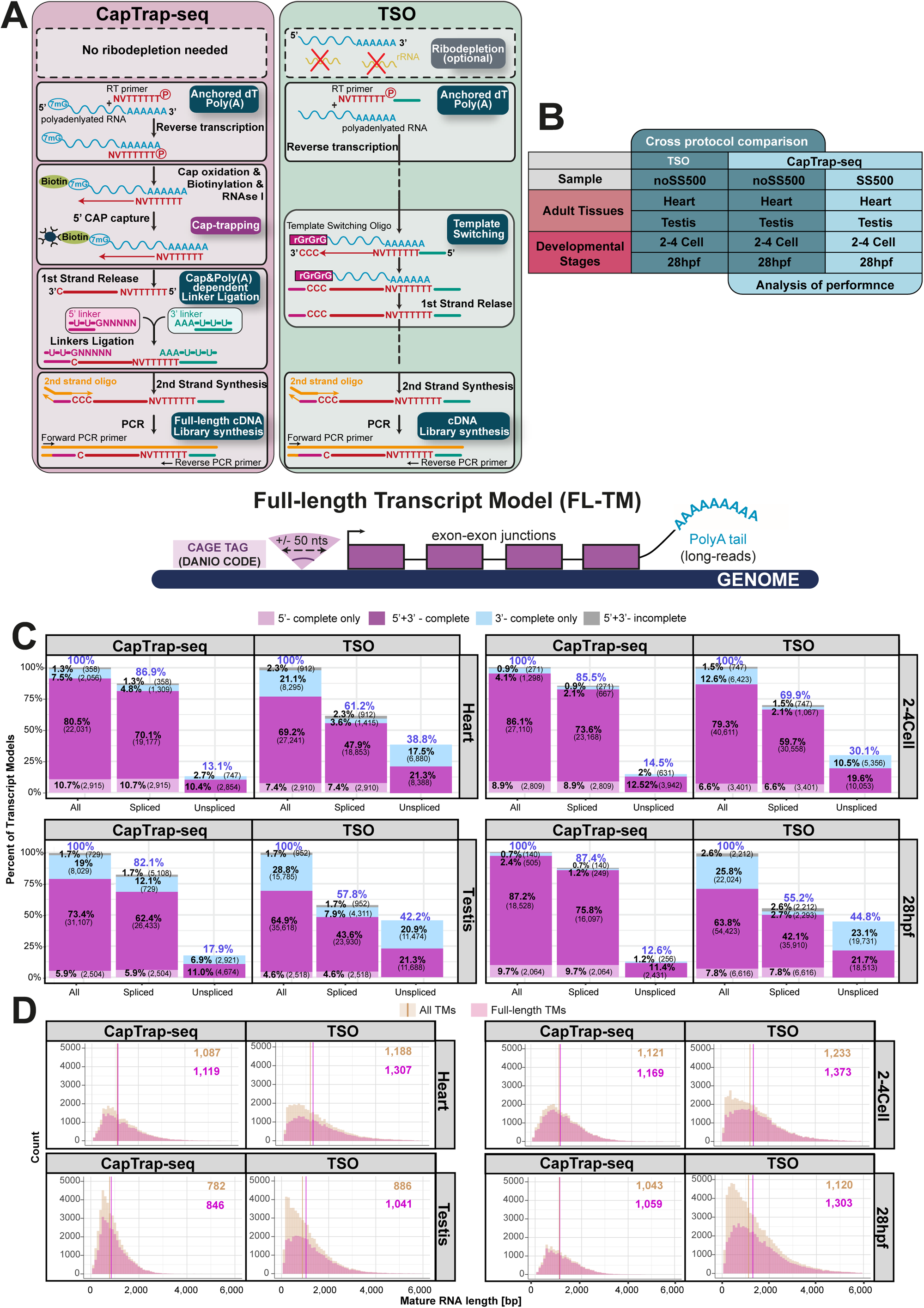
Full-length transcript annotation. **(A)** Comparison of CapTrap-seq and Template Switching Oligo (TSO) Experimental Workflows. Gray sections highlight key steps for each library construction, including Anchored dT Poly(A)+, CAP-trapping^71–74^, CAP and Poly(A)-dependent linker ligation, and FL cDNA library enrichment for CapTrap-seq as described previously^34^, and cDNA library synthesis for the TSO protocol; **(B)** Two samples from adult zebrafish, heart and testis, along with two developmental stages, 2-4 cell and 28 hpf, were selected for a cross-protocol comparison to assess CapTrap-seq performance. The darker blue columns represent comparisons between CapTrap-seq and the TSO protocol, while the lighter blue columns illustrate the performance comparison between the unbiased (CapTrap-seq) and size-selected (CapTrap-seq SS500) samples; (**C)** Identification of full-length transcript models (FL-TMs) among all, spliced and unspliced TMs, based on support from 5′ and 3′ termini using DANIO CODE CAGE tags (N = 1,585,796) and polyA tails from ONT RNA-seq. Colors denote four categories of transcript model completeness: grey represents incomplete TMs, sky blue indicates 3′-complete TMs, light pink represents 5′-complete TMs, and purple shows full-length TMs; **(D)** Length distribution of complete (pink) and incomplete (beige) transcript models for CapTrap-seq and TSO approaches. The median transcript model lengths are indicated in the top right corner, with color coding corresponding to the legend.

Among the protocols tested, CapTrap-seq produced the most uniform read coverage across the bodies of Ensembl-annotated transcripts (Supplementary Figure S1A). Both CapTrap-seq and TSO generated a high and comparable total number of reads per sample, with mapping rates reaching up to 99% (Supplementary Figure S1B). In adult tissues, CapTrap-seq demonstrated lower sequencing error rates than TSO, while in developmental samples, error rates were comparable between the two methods (Supplementary Figure S1C). CapTrap-seq also yielded the highest proportion of complete transcript molecules, with up to 87% of reconstructed transcripts being full-length (Figure 1C). This proportion was notably higher in developmental samples, likely due to the greater availability of DANIO-CODE CAGE data at these stages^31,32^. On average, CapTrap-seq produced shorter reads (Supplementary Figure S1B) and transcript models (Figure 1D) than TSO. This trend held across all reads and TM types, including spliced subsets (Supplementary Figure S1D), consistent with our earlier observations in human and mouse datasets^34^.

A majority (82.1–87.4%) of transcripts detected in CapTrap-seq samples were spliced, compared to 55.2–61.2% in TSO libraries (Figure 1C). Read-level analysis, including High-Confidence Genome Mappings (HCGMs), revealed that CapTrap-seq generates more spliced reads than TSO. Spliced HCGMs are characterized by canonical splice junctions and poly(A) tails, while unspliced HCGMs are defined by the presence of a poly(A) tail. These HCGMs, identified by LyRic^38,39^, filter out poor-quality alignments to infer accurate transcript models. CapTrap-seq consistently produced more spliced HCGMs (54–61%) than TSO (38–41%) (Supplementary Figure S2A). While unspliced reads can result from fragmented transcripts or genomic DNA contamination, the detection of poly(A) tails in unspliced HCGMs mitigates this concern.

Reads generated by CapTrap-seq and TSO mapped to similar gene biotypes (Ensembl v104) with most reads aligning to protein-coding and lncRNA genes (Supplementary Figure S2B). However, TSO produced a higher proportion of non-exonic reads, particularly in testis (17% for TSO vs. 7.5% for CapTrap-seq), which primarily mapped to intergenic or intronic regions. In terms of nucleotide coverage, CapTrap-seq achieved ∼73-84% coverage within genic regions (exons, CDS and UTRs), whereas TSO showed greater variability across samples, with coverage ranging from 60–79% (Supplementary Figure S2C). CapTrap-seq also produced fewer intronic reads (∼6–11%) compared to TSO (∼7–22%), with the largest difference seen in the 28hpf sample (6.1% for CapTrap-seq vs. 22.2% for TSO). Notably, CapTrap-seq effectively eliminated rRNA reads, which remain uncapped under native conditions (Supplementary Figure S2B), thereby minimizing the need for additional ribodepletion. This is especially beneficial considering the limited effectiveness of existing ribodepletion methods for zebrafish, which present a significant challenge in zebrafish transcriptomic studies.

To evaluate the accuracy of transcript reconstruction and the quantitative capability of CapTrap-seq, we employed two sets of external spike-in controls: the External RNA Controls Consortium (ERCC) spike-ins, designed to mimic the natural dynamic range of RNA expression, and the Spike-In RNA Variants (SIRVs), which capture transcriptomic complexity through diverse isoform structures^40–42^. Since their native forms lack a 5′ cap structure, both ERCCs and SIRVs were enzymatically capped using a previously described method^34,43^ incorporated into the sequencing library preparation process. Both protocols accurately detected the majority of SIRVs end-to-end in adult and developmental samples (Supplementary Figure S3A). However, CapTrap-seq enabled deeper sampling of spike-in controls compared to TSO. While TSO detected a higher number of SIRVs, its detection was broader, including more partially detected controls. In contrast, CapTrap-seq provided more consistent detection, with full-length detection of SIRVs when present and complete absence when not detected. CapTrap-seq also reliably quantified ERCC spike-in expression, showing a strong correlation with their known input concentrations (Supplementary Figure S3B). In comparison, TSO exhibited slightly lower correlation and missed a greater number of ERCC spike-ins. It is important to note the bias in this analysis, as TSO was able to detect both capped and uncapped spike-in controls, whereas CapTrap-seq could only detect capped ones.

Despite the high efficiency of enzymatic capping, it is likely that not all molecules are successfully capped, giving TSO an advantage in this case. We compared the performance of CapTrap-seq and TSO protocols against the Ensembl zebrafish genome annotation to assess their effectiveness in detecting annotated genes. Overall, TSO identified more Ensembl genes than CapTrap-seq (Supplementary Figure S4A). However, this broader detection was largely driven by unspliced transcripts. TSO sampled a wider range of genes but at a lower depth, whereas CapTrap-seq offered deeper coverage of a more focused gene set. This targeted approach enabled CapTrap-seq to detect a higher number of transcript isoforms per gene, including full-length alternatively spliced variants with potential functional relevance (Supplementary Figure S4B), except in developmental samples, where TSO was particularly effective. Both methods also supported the discovery of novel genes by identifying unannotated transcripts in the intergenic regions (Supplementary Figure S4C). CapTrap-seq identified more full-length spliced transcripts in intergenic regions of adult samples, while TSO was more effective in developmental samples, likely due to the presence of longer transcripts in these samples^44^. The highest proportion of novel transcripts (∼14-17%) located in intergenic regions was observed in testis, with the majority being full-length spliced transcripts in CapTrap-seq sample. The detection rate of novel genes in developmental and cardiac-related samples, areas where zebrafish is a widely used model was lower with up to 3% of full-length spliced transcripts detected. This underscores the importance of including a diverse set of tissues in transcriptomic analyses, alongside CapTrap-seq, to expand the limited number of annotated genes in zebrafish (Supplementary Figure S5A) and improve the representation of alternatively spliced transcript isoforms (Supplementary Figure S5B).

Specific examples highlight the ability of CapTrap-seq to reliably detect full-length, spliced transcripts for biologically relevant genes. For instance, the CapTrap-seq expanded the annotation of the *LIM Homeobox 9* (*lhx9*) gene, a transcription factor essential for neuron fate specification and thalamus development^45^. Both CapTrap-seq and TSO accurately annotated the *lhx9* gene and refined the transcription termination site (TTS) for the Ensembl *lhx9* annotation (Figure 2A). The detected isoform is supported by the NCBI annotation. In addition, CapTrap-seq enabled detection of novel transcript isoform with alternative TSS upstream of the NCBI and Ensembl annotated TSS. These new TSSs and TTSs were further validated by CAGE tags, consensus promoters, and 3P-seq peaks from DANIO-CODE^31,32^, confirming their accuracy. CapTrap-seq also improved annotation of the *T-box protein 5* (*tbx5b*) gene, a transcription factor critical for zebrafish heart development and pectoral fin formation^46^. Both TSO and CapTrap-seq accurately identified a novel TTS located far downstream of the ENSEMBL-annotated site, while CapTrap-seq additionally revealed an alternative TSS with heart-specific expression (Figure 2B).

**Figure 2.**
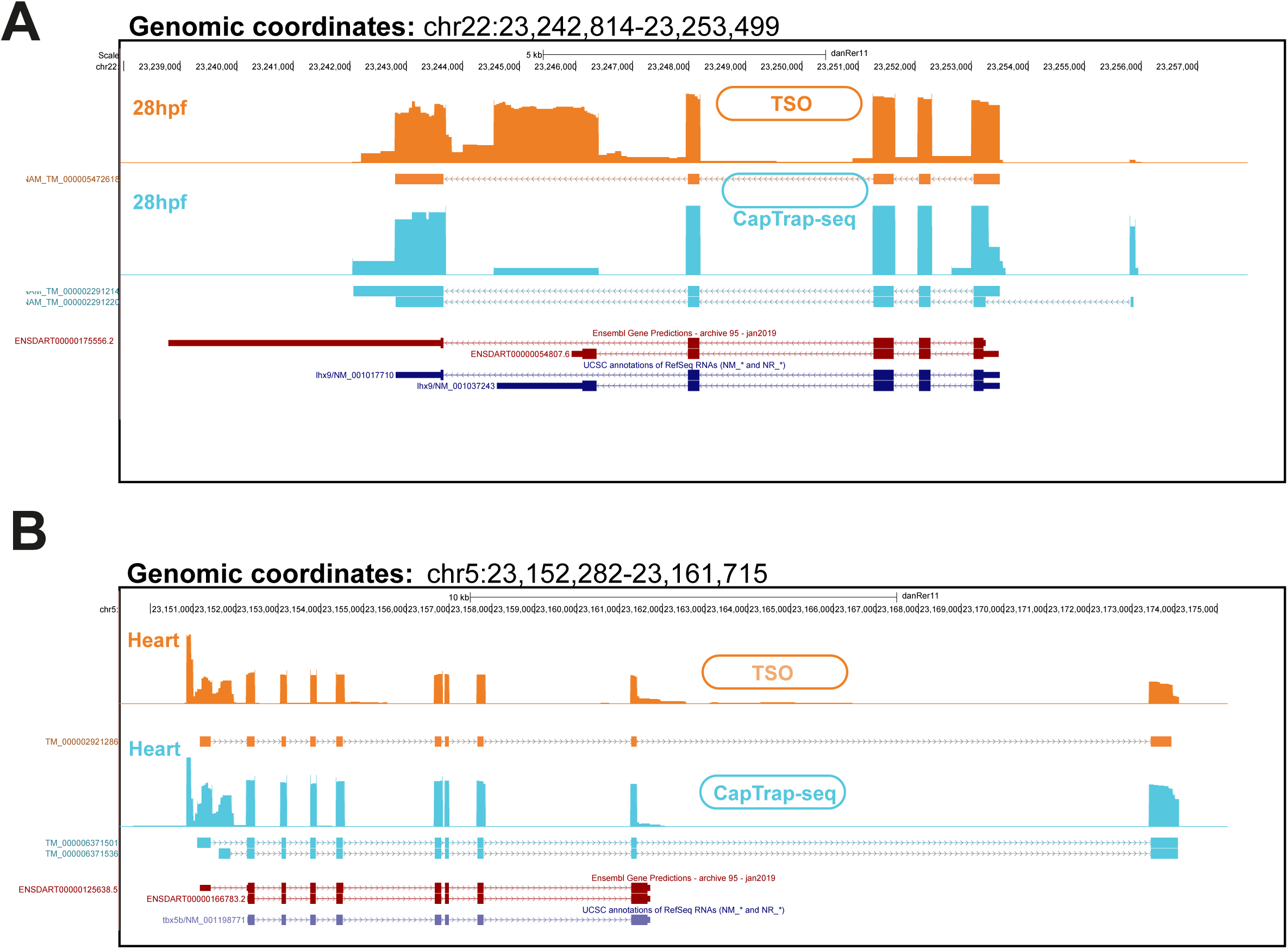
Transcript model reconstruction across library preparation methods. **(A-B)** Transcript models identified for the **(A)** *lhx9* and **(B)** *tbx5b* genes in zebrafish. Colors represent the library preparation method: orange for TSO and blue for CapTrap-seq. Ensembl models are displayed in burgundy, while RefSeq models are shown in navy blue. The bigwig files, generated from long-read ONT RNA-seq data, are shown above each transcript model in full view using the signal track mode on the UCSC Genome Browser. 5′ end support from CAGE seq signals and consensus promoters and 3′ end support from 3P seq data, all sourced from DANIO-CODE^31^, are indicated by vertical black lines.

### Improving CapTrap-seq performance with the size selection process

To improve CapTrap-seq performance and address the reduced detection efficiency of longer transcripts, we added a size selection step. RNA fragments shorter than 500 nucleotides were excluded, enriching the sample for longer transcripts while still allowing detection of RNA molecules across a range of lengths (Supplementary Figure S5C). The 500-nucleotide threshold also aligns with the current minimum length criterion for lncRNA transcripts^47^. Recognizing that size selection can introduce bias in the length distribution of different RNA types^48,49^, we directly compared the performance of size-selected CapTrap-seq samples with their unbiased counterparts (Figure 1B). We observed enhanced read coverage of Ensembl-annotated transcripts after implementing the size selection step (Supplementary Figure 5D). At the read level, size selection increased the median read length in the 28 hpf and testis samples, with only a minimal decrease in the 2–4 cell and heart samples (Supplementary Figure 5E). Interestingly, size selection led to a slight increase in the length of LyRic-inferred TMs (Figure 3A), particularly for spliced TMs (Supplementary Figure S5F). While the overall proportion of full-length transcripts remained consistent between size-selected and unbiased CapTrap-seq samples, the size-selected samples yielded more complete TMs in terms of absolute numbers, except for the heart sample (Figure 3B). Size selection had minimal impact on read accuracy (Supplementary Figure 6A), but it improved polyA tail detection, particularly in developmental stages (Supplementary Figure 6B). No major differences were observed in biotype detection or rRNA removal efficiency following size selection (Supplementary Figure 6C). However, we did observe a slight increase in the detection of novel transcripts in the intergenic space (Supplementary Figure 6D), including full-length spliced isoforms (Supplementary Figure 7A). The improved performance of CapTrap-seq following the inclusion of the size-selection step is evident in specific cases. For example, in the *rab8a* gene (spliced length ∼2kb), size-selected CapTrap-seq enabled the identification of multiple novel isoforms and terminal exons that were not detected using the standard CapTrap-seq protocol (Figure 3C). This modification also increased the overall detection of novel genes and expanded the diversity of alternative isoforms (Supplementary Figure 7B).

**Figure 3.**
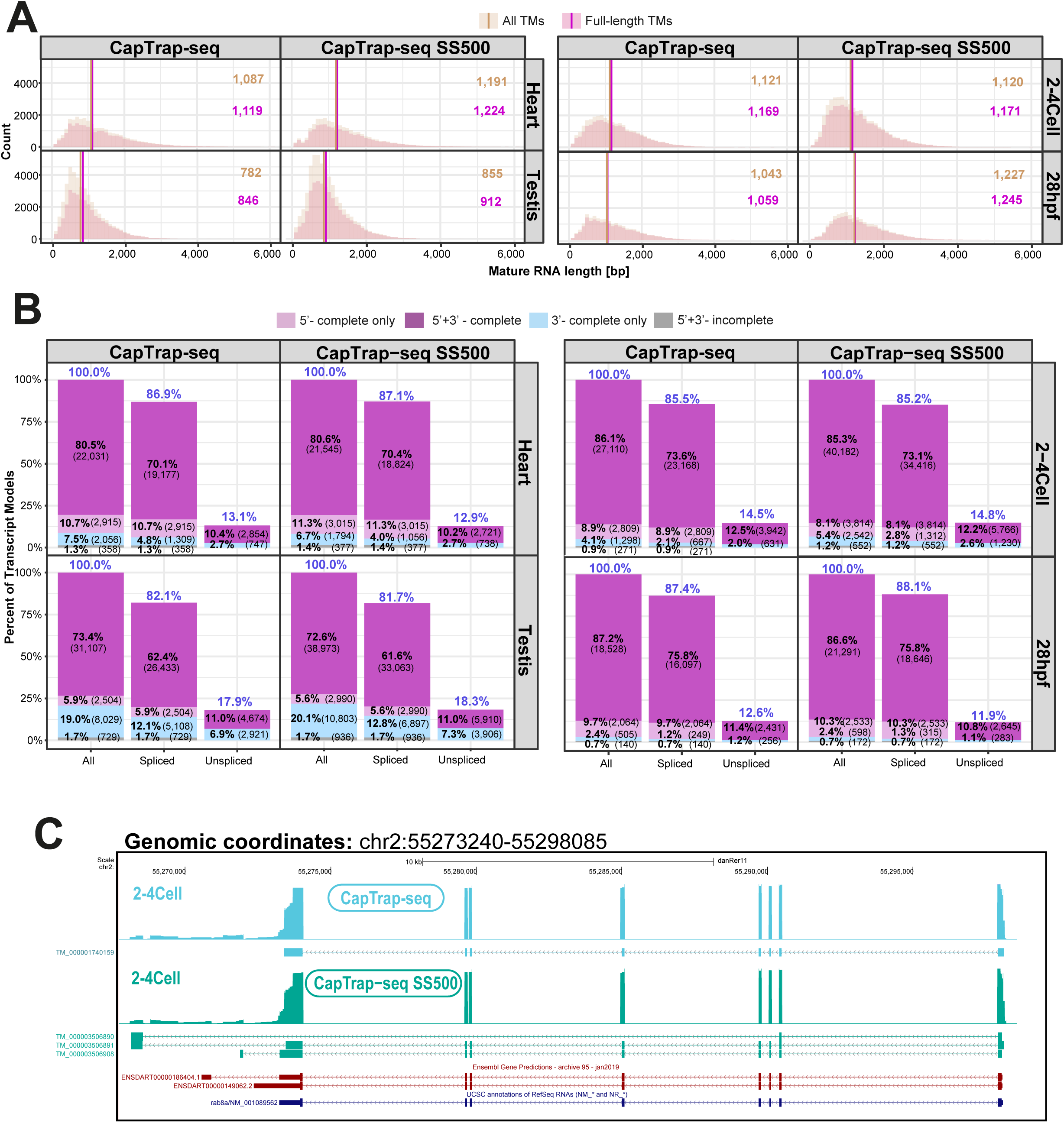
The effect of size-selection on CapTrap-seq performance. **(A)** Length distribution of complete (pink) and incomplete (beige) transcript models for stanard and size-selected (SS500) CapTrap-seq samples. Median transcript model lengths are shown in the top right corner, with colors corresponding to the legend; **(B)** Proportion of transcript models with various types of termini support, as defined in Figure 1C, for standard and size-selected (SS500) CapTrap-seq; **(C)** Transcript models identified for the *rab8a* gene: standard CapTrap-seq shown in blue, and the size-selected (SS500) CapTrap-seq represented in green. Ensembl models are displayed in burgundy, while RefSeq models are shown in navy blue. The corresponding bigwig files from long-read ONT RNA-seq data are displayed above each transcript model in signal track mode, using the full view on the UCSC Genome Browser. Vertical black lines indicate 5′ end support from DANIO-CODE CAGE seq signals and consensus promoters and 3′ end support from DANIO-CODE^31^ 3P-seq data.

We also examined the overlap between transcript sets generated by the tested protocols, defining transcripts as identical if they shared the same intron chains, regardless of differences at the 5′ and 3′ ends. There was substantial overlap, with ∼25% of intron chains shared between standard and size-selected CapTrap-seq and TSO libraries (Supplementary Figure 8). The proportion of protocol-specific intron chains remained relatively consistent across samples, except for the 28hpf sample, where TSO detected the highest proportion of unique intron chains (52% across all transcripts and 46.9% among full-length models). The overlap between technologies increased when focusing exclusively on full-length transcript models. Notably, size-selection had minimal effect on SIRV spike-in detection (Supplementary Figure 9A) and did not compromise the quantitative accuracy of CapTrap-seq (Supplementary Figure 9B).

### Full-length RNA sequencing reveals molecules with novel functional potential

Alternative splicing increases transcript diversity and modulates RNA function by selectively including or excluding specific exons^19^. It plays a crucial role in shaping the functional potential of RNA molecules, orchestrating their interactions with other biological molecules like DNA, RNA, and proteins^50^. To uncover these roles, it is essential to determine the complete structure of RNA transcripts, including their exon composition and arrangement. Both the standard and size-selected CapTrap-seq protocols reliably identified novel, full-length RNA transcripts with distinct functional potential from biologically relevant genes, including those encoding RNA-binding proteins. One example is *hnrnpa1a* (*heterogeneous nuclear ribonucleoprotein A1a*), which encodes a multi-domain protein containing two N-terminal RNA recognition motif (RRMs) domains and a C-terminal glycine-rich low-complexity domain (PrLD) that includes an RGG-repeat region and the M9 nuclear localization signal^51–53^. This protein is a key regulator of RNA metabolism, participating in processes such as splicing, RNA stability, transport, and translational control^54,55^. It also plays dynamic regulatory roles across developmental stages, particularly before and after the maternal-to-zygotic transition (MZT)^56^. Before zygotic genome activation, hnRNP A1a regulates the poly(A) tail length and translation efficiency of maternally inherited mRNAs through sequence-specific binding to their 3′ UTRs. After genome activation, its role transitions to the regulation of pre-mRNA splicing and processing of primary *mir-430* transcripts^56^. In non-mammalian vertebrates, only the long isoform of *hnRNPA1* is typically expressed, with constitutive inclusion of exon 7B/8. The alternative skipping of this exon emerged during mammalian evolution through the acquisition of an intronic silencer element (CE6) that disrupts recognition of its 5′ splice site^57,58^. CapTrap-seq, particularly its size-selected variant, uniquely detected alternative *hnrnpa1a* transcript isoforms in 2–4-cell zebrafish embryos (prior to zygotic genome activation) that lacked exons 6-9 (Figure 4A). These exons encode part of the glycine-rich C-terminal domain, which is essential for nuclear–cytoplasmic transport and spliceosome assembly. Their exclusion likely results in a truncated protein with diminished ability to form higher-order complexes and reduced splicing activity, but potentially increased affinity for the 3′ UTRs of maternal transcripts. This suggests that stage-specific splicing in zebrafish serves as a mechanism to temporarily produce an alternative hnRNP A1 isoform, which may play a role in stabilizing maternal mRNA during early development. Notably, this isoform was absent at 28 hpf (after genome activation) and in adult heart samples.

**Figure 4.**
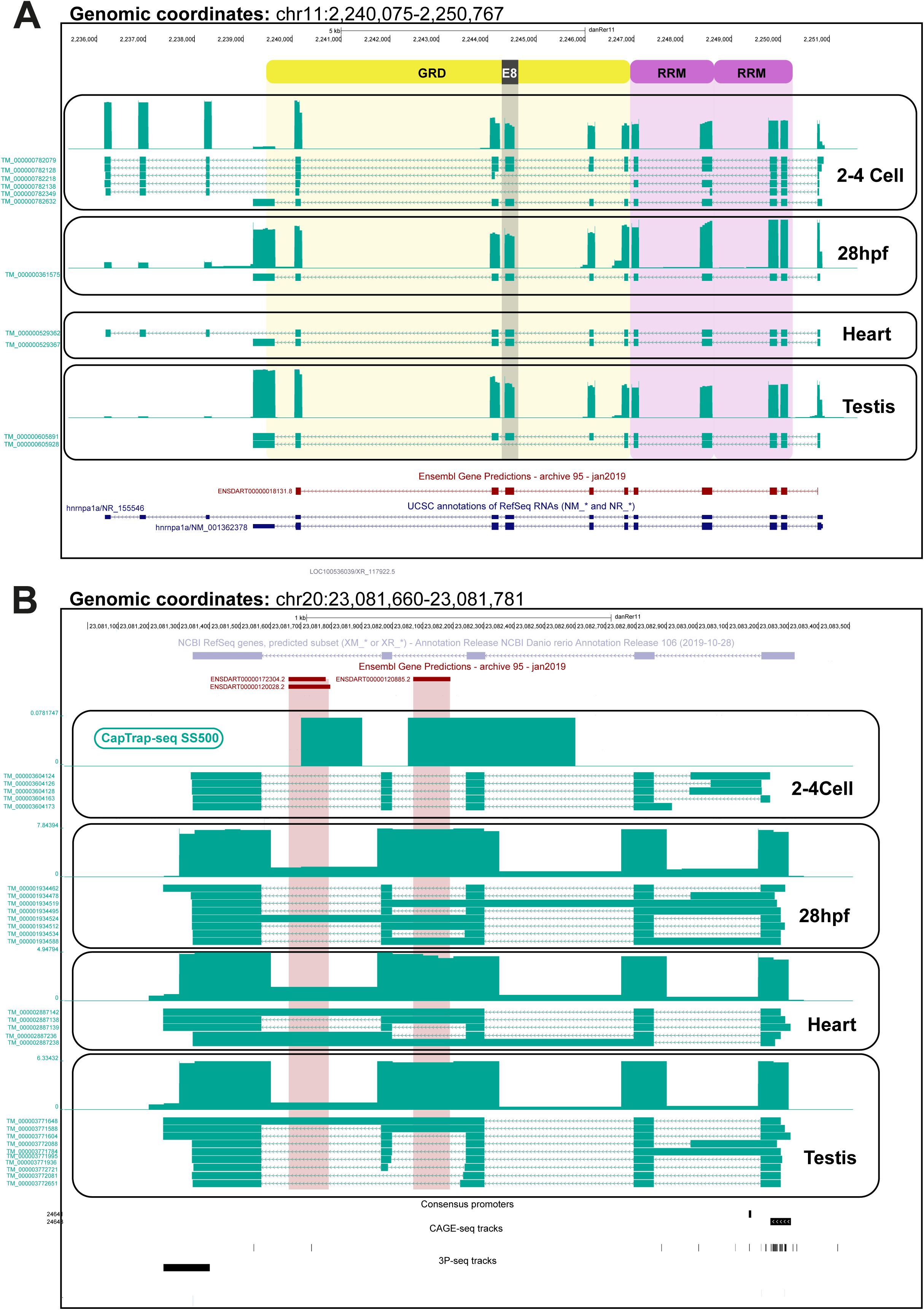
Annotation extension for biologically relevant zebrafish genes. **(A)** Novel transcript isoforms of the *hnrnpa1a* gene detected in size-selected (SS500) CapTrap-seq samples (green) across all biological conditions tested. Exons are colored by its suspected protein-domain contribution: purple for the RNA Recognition Motif (RRM), yellow for the Glycine-Rich Domain (GRD), and gray for exon 8. ENSEMBL models are shown in burgundy and RefSeq models in navy blue. Vertical black lines indicate 5′ end support from DANIO-CODE CAGE-seq signals and consensus promoters, and 3′ end support from DANIO-CODE 3P-seq data. Corresponding bigWig tracks from long-read ONT RNA-seq are displayed above each transcript model in full-view signal-track mode on the UCSC Genome Browser. **(B)** Novel transcript isoforms of the *dancr* lncRNA detected in size-selected (SS500) CapTrap-seq samples (shown in green) across all tested biological conditions. snoRNA gene models annotated by ENSEMBL are shown in burgundy. Vertical black lines mark 5′ end support from DANIO-CODE^31^ CAGE-seq signals and consensus promoters as well as 3′ end support from DANIO-CODE 3P-seq data. Corresponding bigWig tracks from long-read ONT RNA-seq appear above each transcript model in full-view signal-track mode on the UCSC Genome Browser.

In the 2–4-cell stage sample, we also identified an *hnrnpa1a* isoform with a 3′ end extended beyond the Ensembl annotation, corresponding to a transcript cataloged in the NCBI database. This isoform was also detected in adult heart tissue but was absent at 28 hpf and in testes. The functional significance of this alternative transcription termination site, as well as its context-specific role in zebrafish biology, remains to be elucidated.

CapTrap-seq also proved effective in accurately identifying lncRNA molecules, tackling the issue of poor-quality zebrafish lncRNA annotations^30^ (Supplementary Figure 5A-C). Using CapTrap-seq, we identified the *dancr* (*differentiation antagonizing non-protein coding RNA*) lncRNA gene along with its multiple spliced alternative transcript isoforms. *DANCR* is a functional human lncRNA involved in inhibiting cell differentiation^59,60^. It also shows positional conservation in the zebrafish genome^61^. Recent study showed functional significance of *dancr* in zebrafish embryonic development and cell death gene regulation^61^.

Despite its functional importance and expression in multiple distinct cell types, *dancr* has yet to be properly annotated in the latest zebrafish genome assembly (danRer11). While the NCBI gene set predicts *dancr* as an uncharacterized lncRNA locus (LOC100536039), it appears to be entirely absent from the Ensembl annotation, likely due to its computationally derived nature and the lack of manual curation in zebrafish. Our long-read data not only accurately identified *dancr*, but also revealed distinct alternative splicing events. Specifically, intron retention was more frequent at 28 hpf and in heart tissue, while exon skipping was more commonly observed in the testis (Figure 4B). These events were consistently detected across all three library preparation methods (TSO, standard, and size-selected CapTrap-seq), with the most pronounced patterns observed in the size-selected CapTrap-seq samples. The complex splicing landscape of *dancr* mirrors its dynamic expression during zebrafish development and may be linked to its differential subcellular localization^61^. Given its expression in various cell types, alternative transcription start sites may help regulate cell-type-specific isoform expression, while alternative splicing could determine specific subcellular localization. This model, however, requires experimental confirmation. Interestingly, *dancr* contains small nucleolar RNAs (snoRNAs) within its introns, and the majority of intron retention events occurred in snoRNA-containing regions, suggesting a potential regulatory interaction between the host lncRNA and its intronic snoRNAs^62^.

Using CapTrap-seq, we also refined the annotation of *carmn*, a lncRNA known to play a crucial role in heart development by regulating nearby cardiac genes through enhancer-like activity^63,64^, by identifying three novel full-length isoforms, all supported by CAGE and 3P-seq data from DANIO-CODE (Supplementary Figure 10). Notably, two of these isoforms terminate upstream of two annotated miRNA genes, indicating that *carmn* may also have miRNA-independent functions in zebrafish. However, specific validation studies are needed to elucidate such functions.

## DISCUSSION

Zebrafish as a model organism is widely recognized for its genetic and physiological similarities to humans^9,65^, yet their use in large-scale transcriptomic studies lags behind that of mammalian models due to historical reasons and resource-related limitations. In this study, we demonstrate that CapTrap-seq^34^, a genome-agnostic, long-read RNA sequencing method, overcomes key challenges in zebrafish transcriptomics and expands its potential for high-resolution gene expression analysis.

The performance of CapTrap-seq was benchmarked against the widely used Template-Switching Oligo (TSO) method^37^ in both developmental and adult zebrafish samples. This comparison showed that CapTrap-seq enables accurate transcript reconstruction and yields highly reproducible quantitative estimates of transcript abundance, extending its reliability in non-mammalian systems. CapTrap-seq offers significant advantages for transcriptomic research in species beyond humans and mice. First, it efficiently removes uncapped ribosomal RNA without requiring ribodepletion, addressing a major limitation of existing depletion kits in non-mammalian systems. Second, CapTrap-seq enables precise identification of transcription start and termination sites, as well as exon-exon junctions, with single-molecule resolution. This reduces the need for external validation resources like CAGE or 3P-seq data, which are still limited in species like zebrafish^31^. Finally, CapTrap-seq enhances isoform discovery, uncovering previously unknown RNA functions and context-specific alternative splicing events that have implications for development and disease^66,67^.

To mitigate the previously reported bias of CapTrap-seq toward shorter transcripts^34^, we implemented a size-selection step excluding RNAs shorter than 500 nucleotides. This improved the recovery of full-length transcripts by enhancing polyA tail detection, without compromising quantitative accuracy. Future implementations may benefit from a higher size cutoff to further enhance long transcript recovery, alongside efforts to reduce input requirements and polyA+ bias. Additionally, including a diverse range of tissues in transcriptomic studies is essential for expanding gene annotations and discovering novel transcripts.

In summary, CapTrap-seq offers a robust platform for generating high-quality, full-length transcriptomes in zebrafish and other under-annotated species, minimizing the need for extensive manual curation. Its application can help close the annotation gap between zebrafish and mammalian models, enabling broader use of this versatile organism in functional genomics.

## METHODS

### Maintenance of zebrafish lines and Ethical statement

Zebrafish wild-type lines from the AB background were maintained at the zebrafish core facility of the International Institute of Molecular and Cell Biology in Warsaw, Poland (License No. PL14656251), in accordance with institutional and national ethical guidelines for animal welfare. All zebrafish-related procedures were conducted in compliance with ethical principles and regulations.

### Biological sample collection

For organ dissection, adult AB wild-type zebrafish (older than 3 months) were first anesthetized and then euthanized using Tricaine methanesulfonate (MS-222) (Sigma Aldrich, Catalog No. A5040-25G) solution. Heart and testis samples were collected under RNase-free conditions, washed trice in sterile 1x PBS, and placed in 750 μl of TRIzol™ Reagent (Invitrogen, Catalog No. 15596026). Zebrafish embryos were obtained from group spawning of adult AB wild-type individuales which were then incubated at 28°C in egg water (1.5 ml sea salt stock/L distilled water, 60 μg/ml) until desired developmental stage, 2–4 cell stage or 28 hpf. Fifty embryos per stage were collected in an Eppendorf tube, washed three times with sterile 1x PBS buffer, and placed in 1 ml of TRIzol™ Reagent (Invitrogen, Catalog No. 15596026). Tissue and embryo samples in TRIzol™ Reagent were homogenized for 20 s using a Rotor-Stator Tissue Homogenizer (Omni International). The homogenates were then centrifuged (5 min, 12,000 × g, 4°C), and the clear supernatant was incubated for 5 min at room temperature, frozen in liquid nitrogen, and stored at -80°C until RNA isolation.

### RNA samples preparation

Biological samples stored in TRIzol™ Reagent at -80°C were thawed, equilibrated to room temperature, and processed for RNA extraction following the TRIzol™ Reagent Manual with modifications: 1 μl of RNase-free GlycoBlue™ Coprecipitant (15 mg/ml, Invitrogen, Catalog No. AM9515) was added to the aqueous phase for low RNA yield samples, centrifugation after isopropanol precipitation was increased to 20,000 × g for 15 min at 4°C, and RNA pellets were washed twice in 75% ethanol with 10 min centrifugation at 20,000 × g at 4°C between washes. Air-dried RNA pellets were then dissolved in an appropriate volume of RNase-free water. Any residual genomic DNA was removed by incubating the solution with RNase-Free DNase I (QIAGEN, Catalog No. 79254) at room temperature for 10 minutes, following the manufacturer’s guidelines. The RNA samples were subsequently cleaned using Agencourt RNAClean XP beads (Beckman Coulter, Catalog No. A63987) according to the 1.8× reaction volume protocol and eluted in RNase-Free Distilled Water. The total RNA samples were quality-controlled for concentration and integrity. RNA samples (RIN ≥ 8.5) were pooled from 1,000 embryos per developmental stage, 50 adults (25 females, 25 males) for heart, and 50 males for testis. The pooled RNA was mixed with 0.1 volume of 3M RNase-free sodium acetate (pH 5.5) and 2.5 volumes of ice-cold 100% ethanol, then precipitated overnight at -20°C. Pellets were collected by centrifugation (18,000 × g, 30 min, 4°C), washed twice with 75% ethanol, air-dried, and resuspended in nuclease-free water. RNA samples were checked for genomic DNA contamination by PCR amplification of a region of the zebrafish *eef1a1a* gene containing an intron (GRCz11; chr13: 27,317,027-27,317,204; forward strand), which is spliced out in processed RNA. Amplicons from the PCR reaction differed in length: 93 bp for cDNA and 178 bp for genomic DNA.

### cDNA sequencing library preparation

Four total RNA samples, derived from adult heart, testis, and two developmental stages (2–4 cell stage and 28 hpf), were used to benchmark two long-read library preparation methods:

### CapTrap-seq full-length cDNA libraries

5’-capping of external spike-ins (SIRV-Set 4 (Iso Mix E0 / ERCC / Long SIRVs, Lexogen, Catalog No. 141.03)) was performed following a previously described protocol^43^.

CapTrap-seq cDNA libraries were prepared as described^34,35^ with modifications. Specifically, 5 μg total RNA was combined with 3 μl of a 1:100-diluted, pre-5’-capped SIRV-Set 4 (Iso Mix E0 / ERCC / Long SIRVs, Lexogen). UMI16 (100 μM; 5’TCGTCGGCAGCGTCAGATGTGTATAAGAGACAGNNNNNNNNNNNNNNNNGTGGTATCAACGCAGAGTA3’) replaced UMI8 as the second-strand primer. Second-strand cDNA was used directly for LA-PCR with TaKaRa LA Taq (Cat. No. RR002M, Takara) without a drying step. To minimize PCR duplicates, the cDNA was split into two independent PCR reactions, each containing 18.5 μl nuclease-free water, 5 μl 10× LA PCR Buffer II, 8 μl dNTP mix (2.5 mM each), 2.5 μl each of CapTrap_FOR (10 μM; 5’TCGTCGGCAGCGTC3’) and CapTrap_REV (10 μM; 5’GTCTCGTGGGCTCGG3’), 0.5 μl TaKaRa LA Taq (5 U/μl), and 13 μl second-strand product. PCR conditions were: 30 s at 95°C; 16 cycles of 15 s at 95°C, 15 s at 55°C, and 8 min at 68°C; followed by 10 min at 68°C and a 4°C hold. The PCR products were pooled, purified using AMPure XP beads (1× ratio), and resuspended in nuclease-free water. Samples were quantified with Qubit (Qubit 4 Fluorometer, Thermo Fisher Scientific) and quality checked with BioAnalyzer (Agilent 2100 Bioanalyzer, Agilent Technologies).

### Library preparation using Template Switching approach

First, 5 μg of total RNA per sample (Heart, Testis, 2–4 cell, 28 hpf) was mixed with 3 μl of a 1:100-diluted, pre-5’-capped SIRV-Set 4 (Iso Mix E0 / ERCC / Long SIRVs, Lexogen) and undegro rRNA depletion step with the riboPOOL kit for Danio rerio (SiTools, Cat. No. 27DP-K012-000010) following manufacturer’s protocol (riboPOOLKitManual_v6) with modifications: To protect RNA from degradation, 1 μl of RNasin Plus RNase Inhibitor (Promega, Catalog No. N2611) was added to each 13 μl of RNA sample. For optimal hybridization of oligonucleotides to rRNA molecules, the temperature was gradually decreased from 68°C to 37°C at a ramp rate of 0.1°C/s. After rRNA depletion, RNA was cleaned with Agencourt RNAClean XP beads (1.8× ratio), washed three times with 70% ethanol, and eluted in RNase-free water. Samples were quantified with Qubit (Thermo Fisher Scientific). 4 μl of ribo-depleted RNA per reaction was used for first-strand cDNA synthesis with Template Switching RT Enzyme Mix (NEB, Cat. No. M0466), Template Switching Oligo (75 µM; 5’AAGCAGTGGTATCAACGCAGAGTACATrGrG+G3’), and anchored oligo-dT (10 µM; 5’AAGCAGTGGTATCAACGCAGAGTACT30VN3’). Samples were processed in duplicate per biological sample following the manufacturer’s protocol. First-strand cDNA duplicates were pooled, purified with Agencourt RNAClean XP beads (1.8× ratio), eluted in 40 μl RNase-free water, and subjected to second-strand synthesis and amplification with TaKaRa LA Taq (Cat. No. RR002M, Takara). To minimize PCR duplicates, the cDNA was split into two independent PCR reactions, each containing 14 μl nuclease-free water, 5 μl 10× LA PCR Buffer II, 8 μl dNTP mix (2.5 mM each), 2.5 μl ISPCR oligo (10 μM; 5’AAGCAGTGGTATCAACGCAGAGT3’), 0.5 μl TaKaRa LA Taq, and 20 μl second-strand product. PCR conditions were: 30 s at 95°C; 9 cycles of 15 s at 95°C, 15 s at 65°C, and 8 min at 68°C; followed by 10 min at 68°C and a 4°C hold. PCR replicates were pooled, purified with AMPure XP beads (1× ratio), and resuspended in RNase-free water. Samples were quantified using the Qubit 4 Fluorometer (Thermo Fisher Scientific) and quality-checked with the Agilent 2100 Bioanalyzer (Agilent Technologies).

### cDNA size selection

The CapTrap-seq cDNA library was size-selected using the ProNex® Size-Selective Purification System (Promega, Catalog No. NG2001) to enrich cDNA fragments longer than 500 bp. 2 μg of the cDNA sample were mixed with ProNex beads at a 1:1.1 ratio and incubated at room temperature for 10 minutes on the HulaMixer™. Bead-bound cDNA fragments longer than 500 bp were separated using a magnetic rack, washed twice with 80% ethanol, air-dried, and eluted in 20 μl of Elution Buffer at 37°C for 10 minutes with gentle rotation. The supernatant fraction, containing shorter cDNA fragments, was purified by adding 2.0× volume of AMPure XP Reagent (Beckman Coulter, Catalog No. A63881), incubating at room temperature for 10 minutes on the HulaMixer™, washing twice with 80% ethanol, air-drying, and eluting in 20 μl of Nuclease-free water at 37°C for 10 minutes with gentle rotation. Both fractions were assessed for quality using the Agilent 2100 Bioanalyzer and quantified with a Qubit fluorometer.

### ONT sequencing

Samples for Oxford Nanopore Technologies (ONT) MinION long-read sequencing were prepared from TSO, CapTrap-seq, and CapTrap-seq SS500bp libraries using the Nanopore Ligation Sequencing Kit (SQK-LSK109), following the manufacturer’s protocol (version ACDE_9064_v109_revG_23May2018). To achieve optimal loading concentrations for MinION sequencing, 500 ng of each cDNA library was used as input material. 150 fmol of each Nanopore sequencing library was loaded onto separate R.9.4.1 flow cells (FLO-MIN106D) and sequenced for at least 72 hours.

### Data analysis with the LyRic pipeline

Oxford Nanopore Technologies long-read sequencing data generated from both CapTrap-seq and TSO protocols were processed using LyRic, a flexible and automated transcriptome annotation and analysis workflow^39^. Reads were aligned to the zebrafish reference genome (GRCz11/danRer11), supplemented with 96 ERCC and 69 SIRV spike-in controls, using Minimap2^68,69^. A custom reference annotation was created by combining Ensembl gene annotation (v104), SIRV, and ERCC datasets.

Read coverage profiles along annotated Ensembl genes were generated using the depTools2 package^70^, specifically employing the computeMatrix and plotProfile functions. High Confidence Genome Mappings (HCGMs) were defined as read alignments that: (1) contained only canonical splice junctions (spliced reads only), (2) showed no signs of reverse transcription artifacts (spliced reads only), (3) minimum average sequencing quality around the splice junctions (spliced reads only), and (4) included a polyA tail (for spliced and unspliced reads). HCGMs were further merged into non-redundant transcript models using the tmerge function integrated within LyRic. The resulting TMs were merged with the Ensembl (v104) annotation, and novel loci were identified and classified as either intronic or intergenic using LyRic’s buildLoci tool. Comprehensive workflow documentation is available at the LyRic repository^39^, with data and configuration files accessible via GitHub.

## Data availability

All sequencing data, including raw ONT long-read RNA-seq, are available in the GEO repository under accession number GSE294116. Source data are provided in this paper and can be accessed at https://github.com/cobRNA/CapTrap-seq_Zebrafish.git.

## Code availability

The LyRic code is available at https://github.com/guigolab/LyRic^39^. Additionally, the custom code for processing the raw data can be found at https://github.com/cobRNA/CapTrap-seq_Zebrafish.git.

## Acknowledgements

We are grateful to the Guigó laboratory for their valuable contributions and assistance with sample handling. Special thanks go to Carme Arnan Ros and Emilio Palumbo for their support. We also thank Kuznicki’s lab (IIMCB) for both laboratory and scientific assistance. The IIMCB Zebrafish Core Facility is acknowledged for providing services and fish material. This work, including its publication, was supported by the National Science Center, Poland (grant no. 2018/31/B/NZ2/01940 awarded to B.U.-R. and grant no. 2022/45/N/NZ2/03622 awarded to M.K.). S.C-S was supported by Award Number 2U24HG007234-09 to R.G. from the National Human Genome Research Institute. The content of this manuscript is solely the responsibility of the authors and does not necessarily represent the official views of the National Human Genome Research Institute or the National Institutes of Health. We acknowledge support of the Spanish Ministry of Science and Innovation through the Centro de Excelencia Severo Ochoa (CEX2020-001049-S, MCIN/AEI /10.13039/501100011033), and the Generalitat de Catalunya through the CERCA programme. We also extend our thanks to Katarzyna Solka (IBCH PAS) and Romina Garrido (CRG) for her administrative support.

## Author contributions

B.U.-R., and M.K. designed the experiment. M.K. optimized the CapTrap-seq for long-read sequencing with the input from S.C.-S and R.G., generated cDNA libraries and performed the cDNA sequencing. B.U.-R., M.B., and T.M. analyzed the data. B.U-R. wrote the manuscript with contributions from M.K, S.C.-S. and R.G.

---

**Figure S1.**
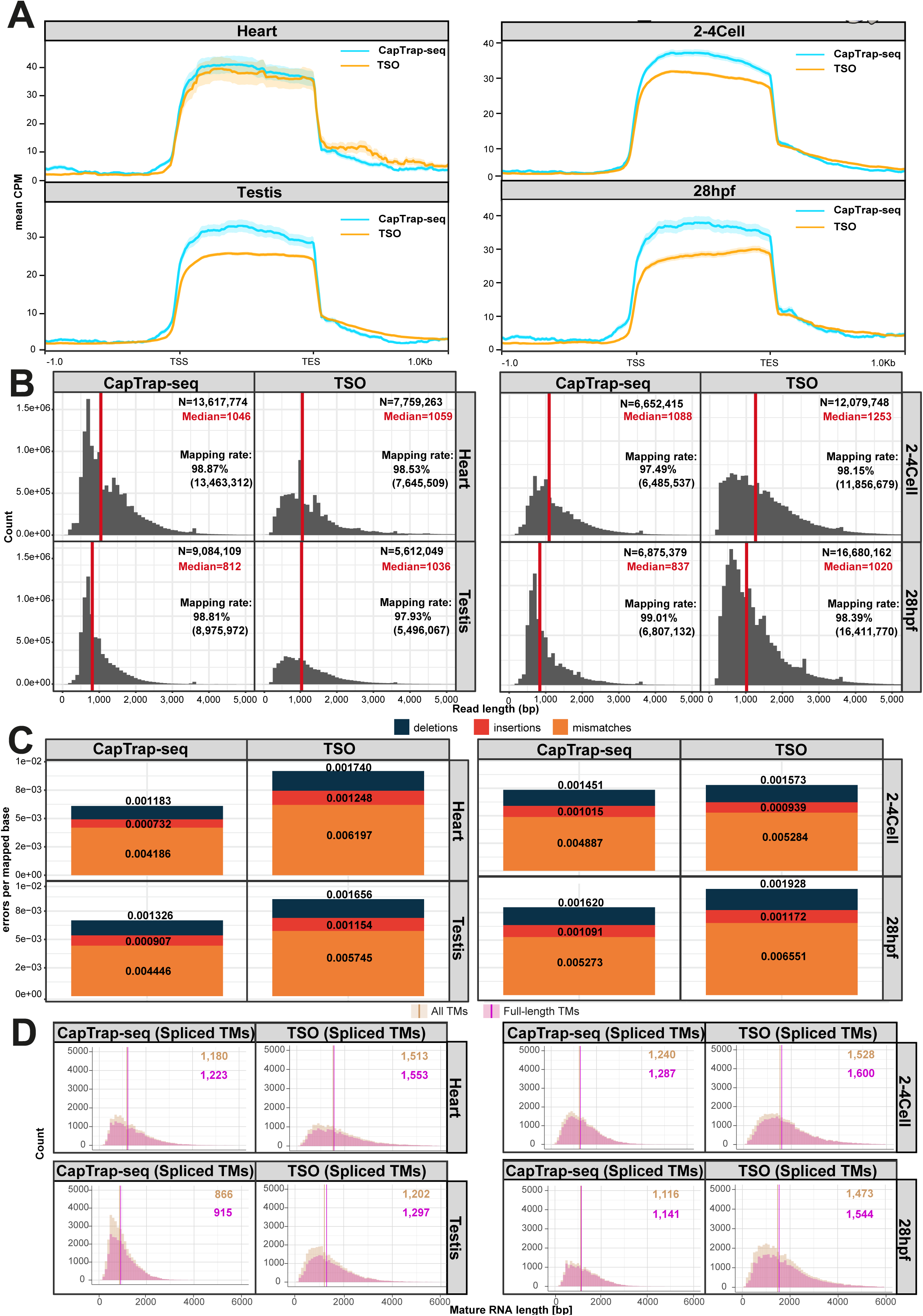
Quality assessment of reads and transcript models across library preparation protocols. **(A)** Read aggregate profiles generated with deepTools^70^ across the bodies of Ensembl-annotated zebrafish genes. CapTrap-seq is shown in blue, and TSO is shown in orange; **(B)** Length distribution of raw long-read ONT reads for each protocol across biological samples. The total number of reads (N), median read length (marked by a red vertical line), and mapping rate are shown in the top right corner of each panel; **(C)** Error rates observed across CapTrap-seq and TSO library preparation protocols for zebrafish samples. Error types are color-coded: deletions (black), insertions (red), and mismatches (orange); **(D)** Length distribution of complete (pink) and incomplete (beige) spliced transcript models generated by CapTrap-seq and TSO approaches. Median transcript model lengths are shown in the top right corner, with colors corresponding to the legend.

**Figure S2.**
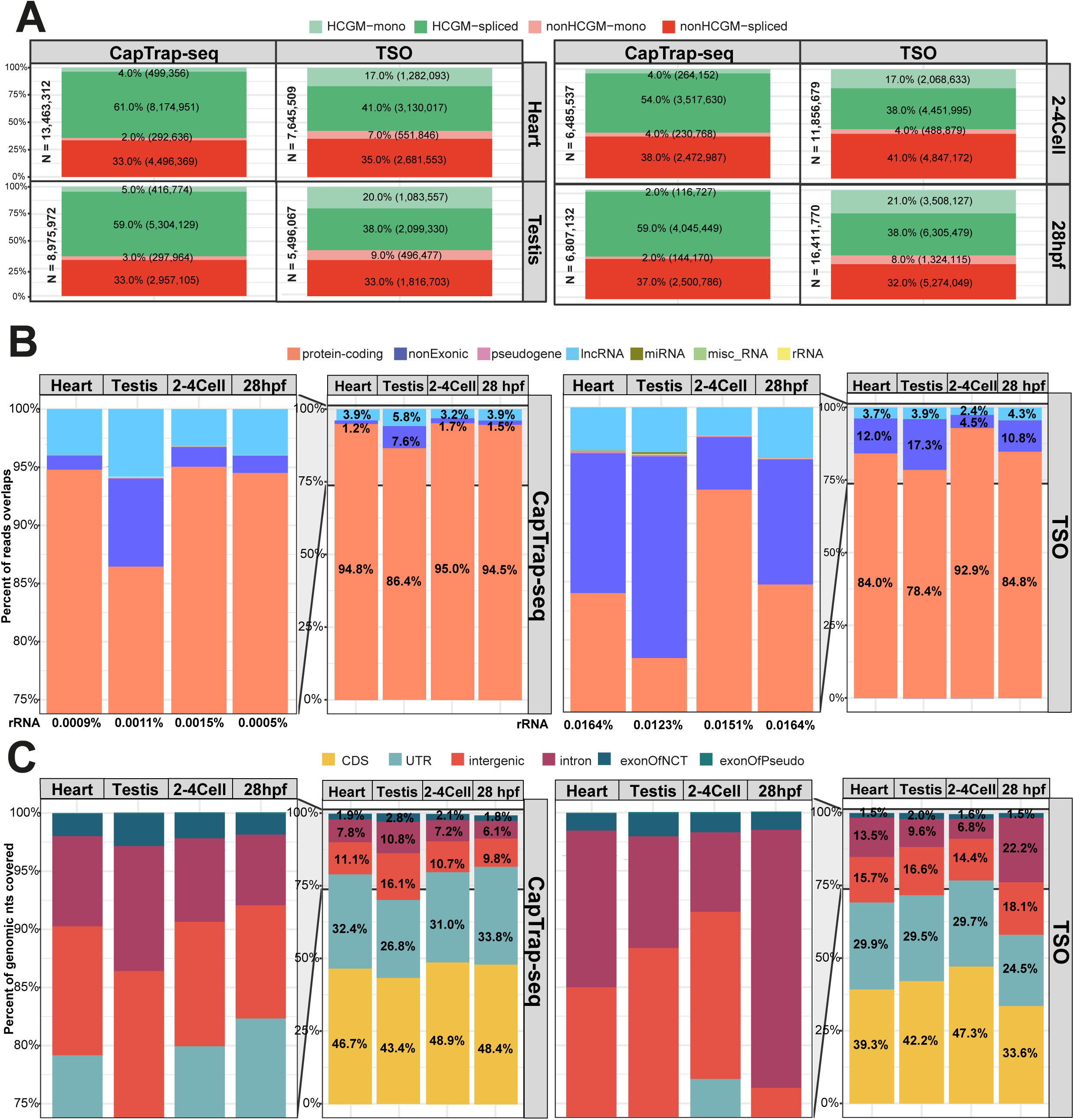
Analysis of read quality across different library preparation protocols. **(A)** The proportion of detected High-Confidence Genome Mappings (HCGMs). The four main classes include: unspliced HCGMs (light green), spliced HCGMs (dark green), unspliced non-HCGMs (light red), and spliced non-HCGMs (dark red). (N) represents the total number of mapped reads; **(B)** Ensembl gene biotype detection by ONT sequencing for CapTrap-seq and TSO protocols. The stacked bar plots show the percentage of raw reads mapping to various gene biotypes represented by different colors: protein-coding (orange), ribosomal RNAs (yellow), pseudogenes (pink), miscellaneous RNAs (green), miRNAs (olive green), non-exonic (dark blue), and lncRNAs (light blue); **(C)** Proportion of nucleotide coverage across various genomic regions. Colors represent specific region types: exon of pseudogene (green), intron (pink), UTR (blue), intergenic (orange), exon of noncoding transcript (navy), and CDS (yellow).

**Figure S3.**
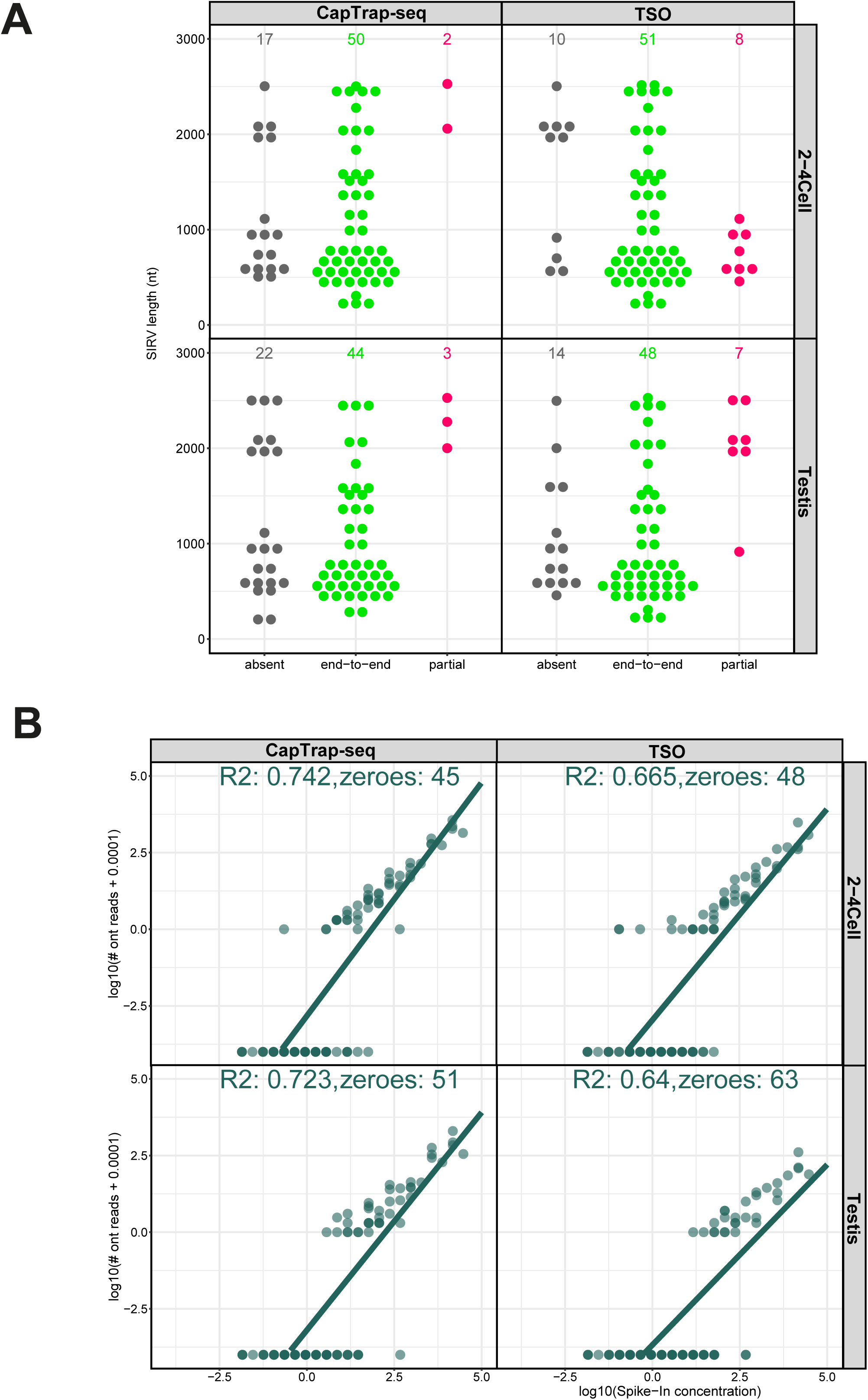
Evaluating CapTrap-seq performance in transcript reconstruction and quantification with RNA spike-in controls. **(A)** Detection of SIRVs relative to transcript length. SIRVs are categorized into three detection levels: end-to-end (green), partial (red), and undetected or absent (gray); **(B)** Correlation between input RNA concentration and raw read counts for ERCC spike-ins in the 2-4 cell and testis samples. Each point represents an individual synthetic ERCC control. The green line indicates a linear regression fit to the dataset, with the corresponding R^2^ value displayed at the top.

**Figure S4.**
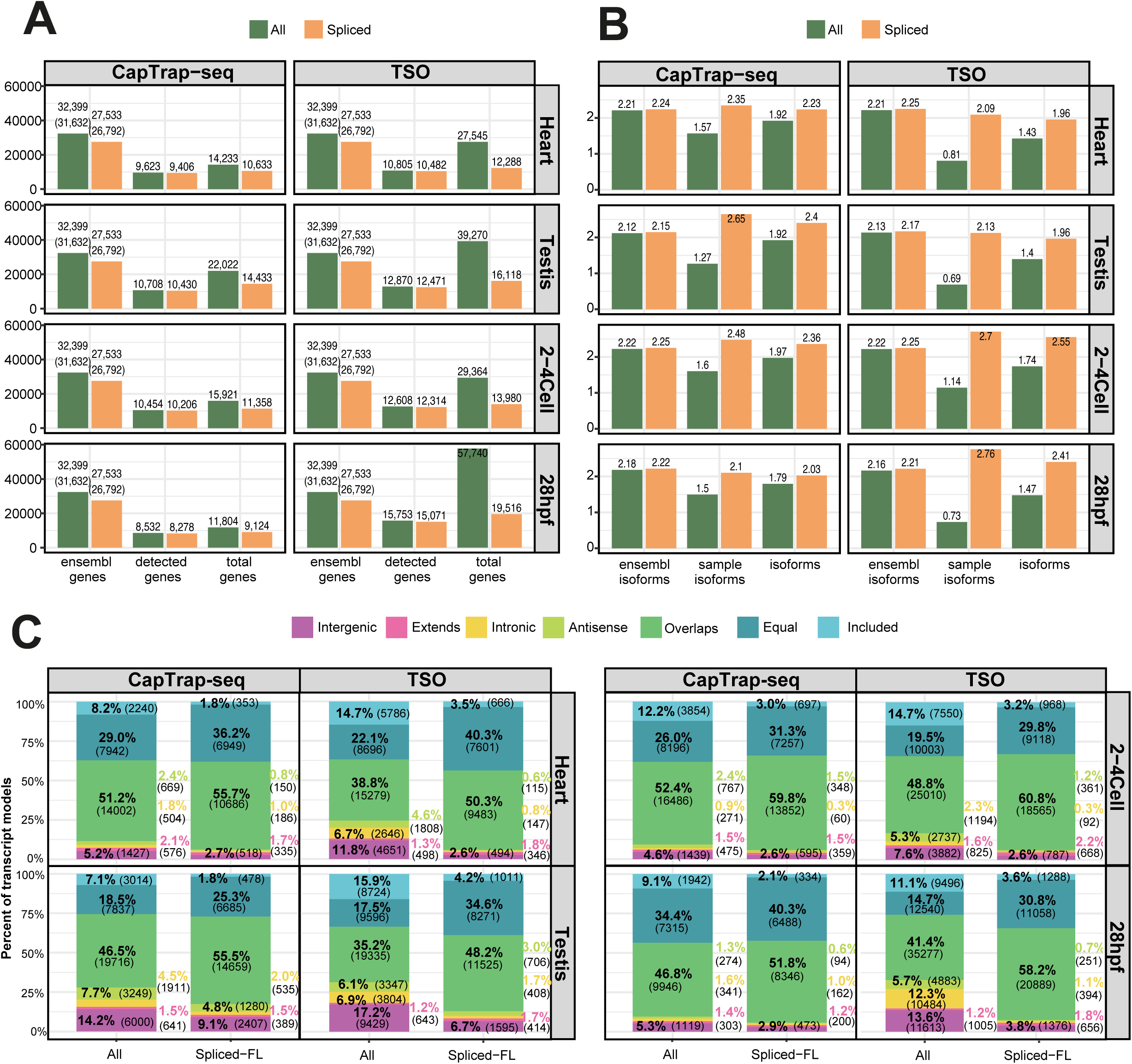
The effect of library preparation protocol on the detection of zebrafish genes. **(A)** Detection rates of Ensembl-annotated and novel genes **(B)** and their alternative isoforms detected by CapTrap-seq and TSO protocols. The "All" and "Spliced" categories are represented in green and orange, respectively; **(C)** Comparison of detected transcript model structures with Ensembl annotation (v104). Transcript models are categorized into seven classes based on their relationship to existing annotations: Included, Equal, Overlaps, Antisense, Intronic, Extends, and Intergenic, each represented by a distinct color code. The classes are simplified versions of gffcompare^75^ transcript classification code.

**Figure S5.**
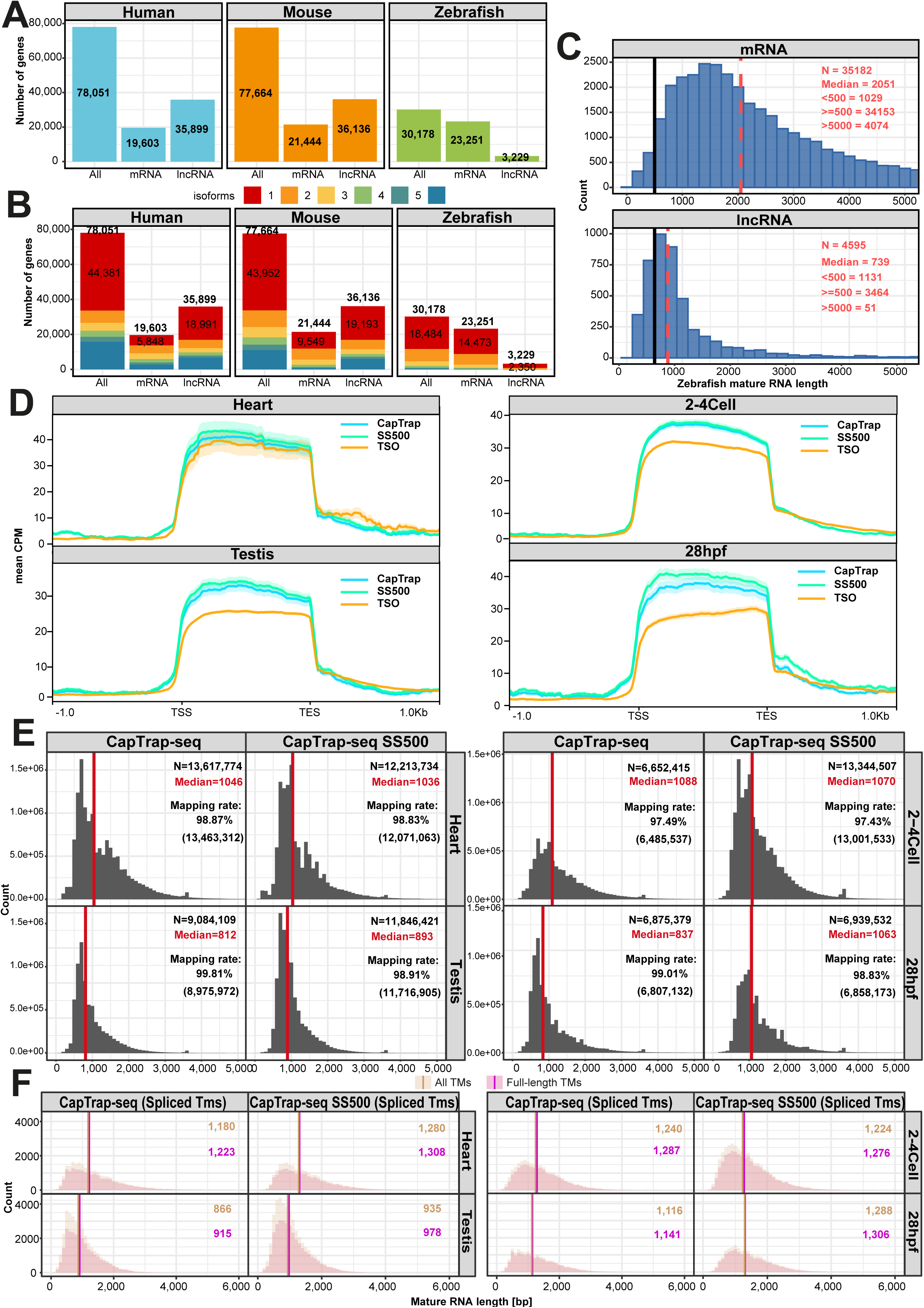
Benchmarking the size-selected CapTrap-seq protocol. **(A)** The number of protein-coding and long non-coding genes annotated in human (shown in blue, GENCODE, v47)^38^, mouse (shown in orange, GENCODE, vM36)^38^, and zebrafish (shown in green, Ensembl, v104); **(B)** The number of isoforms per gene for protein-coding and noncoding biotypes annotated in the human (GENCODE, v47)^36^, mouse (GENCODE, vM36)^36^, and zebrafish (Ensembl, v104) genomes. Genes with varying isoform counts are depicted in different colors; **(C)** Length distribution of zebrafish mRNAs and lncRNAs. The median transcript length is indicated by a red dashed line, while the size-selection cutoff at 500 bp is represented by a black vertical line. The total number of transcripts (N), median length, and counts for transcripts <500 bp, ≥500 bp, and >5000 bp are shown in the top right corner; **(D)** Read aggregate profiles generated with deepTools^70^ across the bodies of Ensembl-annotated zebrafish genes. TSO is shown in orange, unbiased CapTrap-seq in blue, and size-selected sample (SS500) in green; **(E)** The length distribution of raw long-read ONT reads across biological samples, including standard and size-selected (SS500) CapTrap-seq samples . The total number of reads (N), the median read length (indicated by a red vertical line), and the mapping rate are displayed in the top right corner; **(F)** Distribution of lengths for full-length (pink) and incomplete (beige) spliced transcript models in CapTrap-seq and TSO approaches. The median transcript model lengths are shown in the top right corner, with color coding corresponding to the legend.

**Figure S6.**
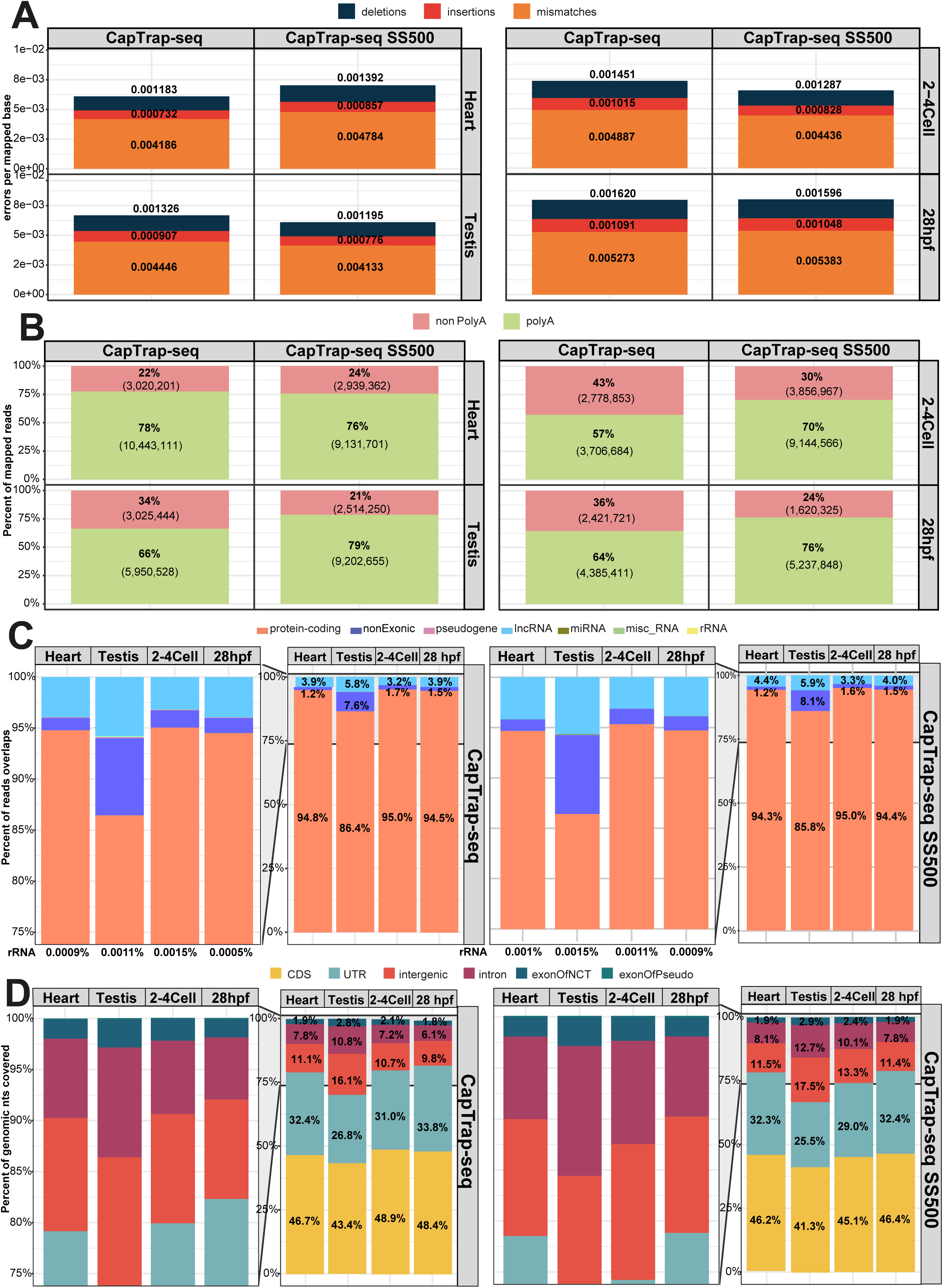
The impact of size-selection on read and transcript model quality. **(A)** Error rates across standard and size-selected (SS500) CapTrap-seq sequencing libraries. Errors are categorized by color: deletions (black), insertions (red), and mismatches (orange); **(B)** Proportion of poly(A) (green) and non-poly(A) (red) ONT reads detected in standard and size-selected (SS500) CapTrap-seq across zebrafish samples; **(C)** Ensembl gene biotype detection by ONT sequencing by both CapTrap-seq protocols. The stacked bar plots show the percentage of raw reads mapping to Ensembl-annotated gene biotypes, with different colors representing: protein-coding (orange), ribosomal RNAs (yellow), pseudogenes (pink), miscellaneous RNAs (green), miRNAs (olive green), non-exonic (dark blue), and lncRNAs (light blue); **(D)** The nucleotide coverage across various genomic regions for standard and size-selected (SS500) CapTrap-seq protocols. Colors represent specific regions: exon of pseudogene (green), intron (pink), UTR (blue), intergenic (orange), exon of noncoding transcript (navy), and CDS (yellow).

**Figure S7.**
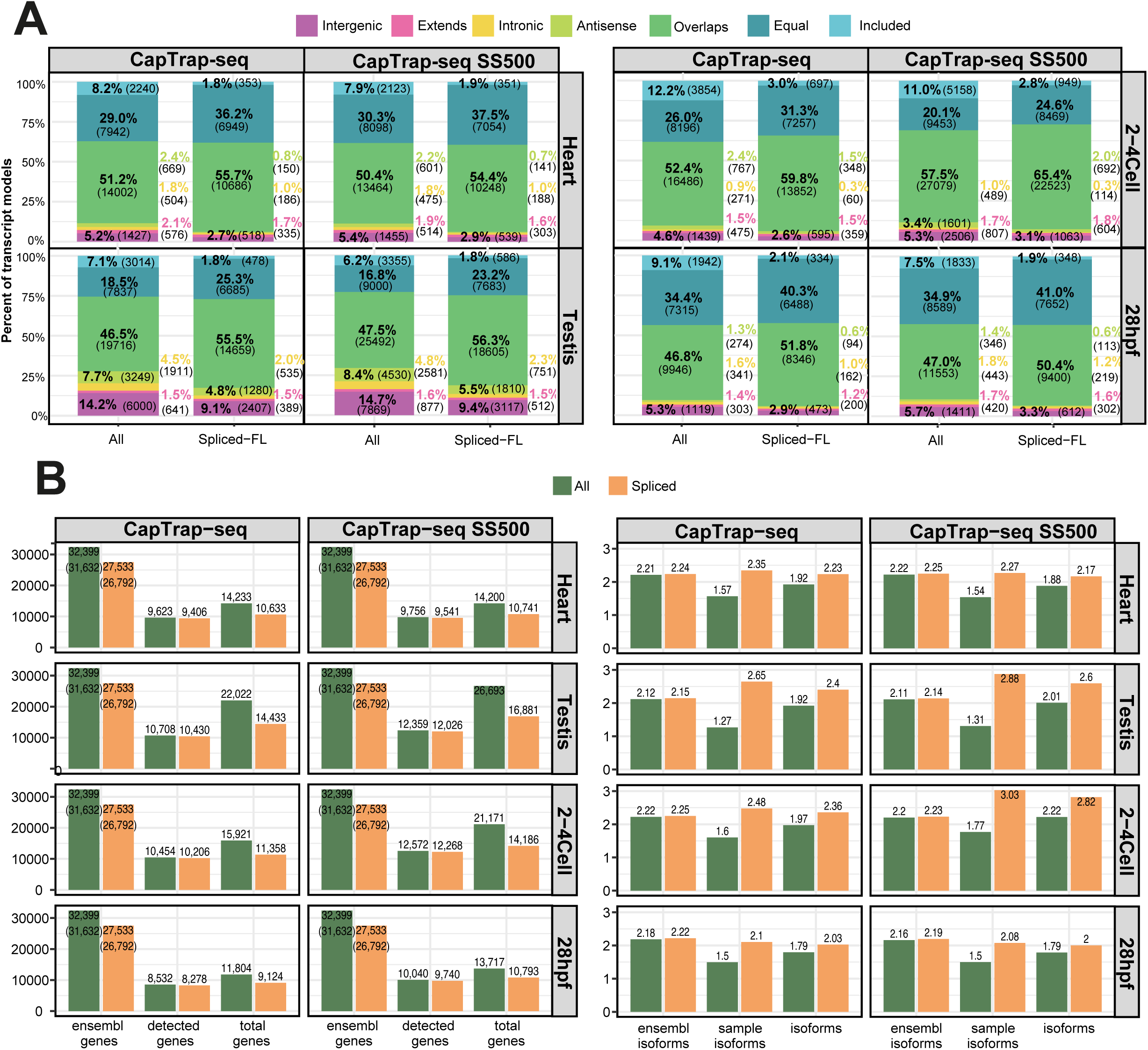
Extending Ensembl annotation with size-selected CapTrap-seq protocol. **(A)** Comparison of transcript model structures generated by standard and size-selected (SS500) CapTrap-seq protocols with respect to the Ensembl annotation reference. Transcript models are categorized into seven classes based on their relationship to existing annotations: Included, Equal, Overlaps, Antisense, Intronic, Extends, and Intergenic, each represented by a distinct color code. The classes are simplified versions of gffcompare^75^ transcript classification code; **(B)** The number of Ensembl-annotated and novel genes and corresponding isoforms for each sample along with genes and isoforms detected by both CapTrap-seq protocols. The "All" and "Spliced" categories are represented in green and orange, respectively.

**Figure S8.**
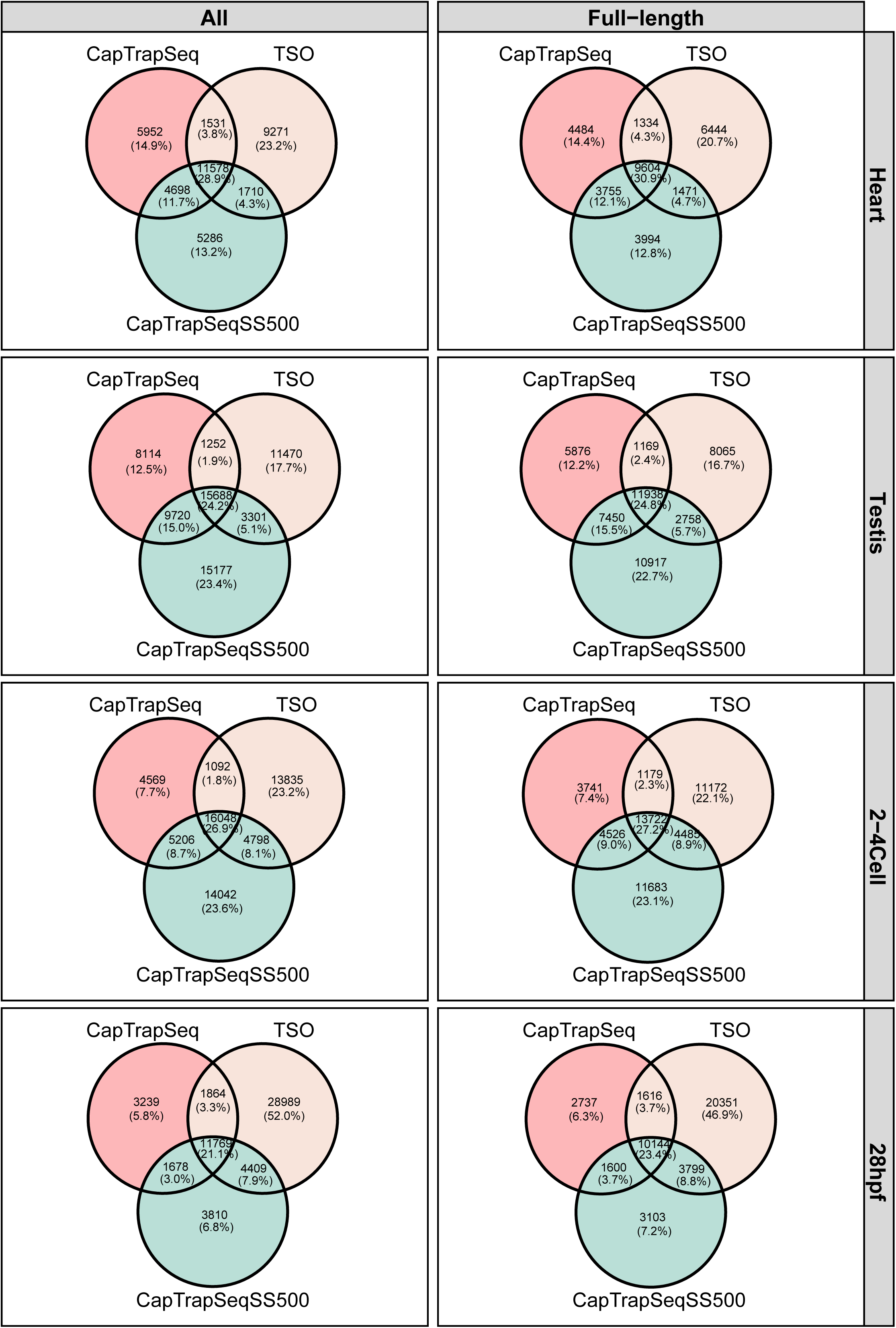
Evaluation of Protocol Specificity. Three-way Venn diagrams illustrating the number of identical transcript models (TMs), both all and full-length, detected by three library preparation methods: standard CapTrap-seq (peach), size-selected CapTrap-seq (green), and TSO (beige). Transcripts were considered identical if they shared the same intron chains, regardless of variations at the 5’ and 3’ ends. Venn diagrams were prepared using the VennDiagram^76^ R package.

**Figure S9.**
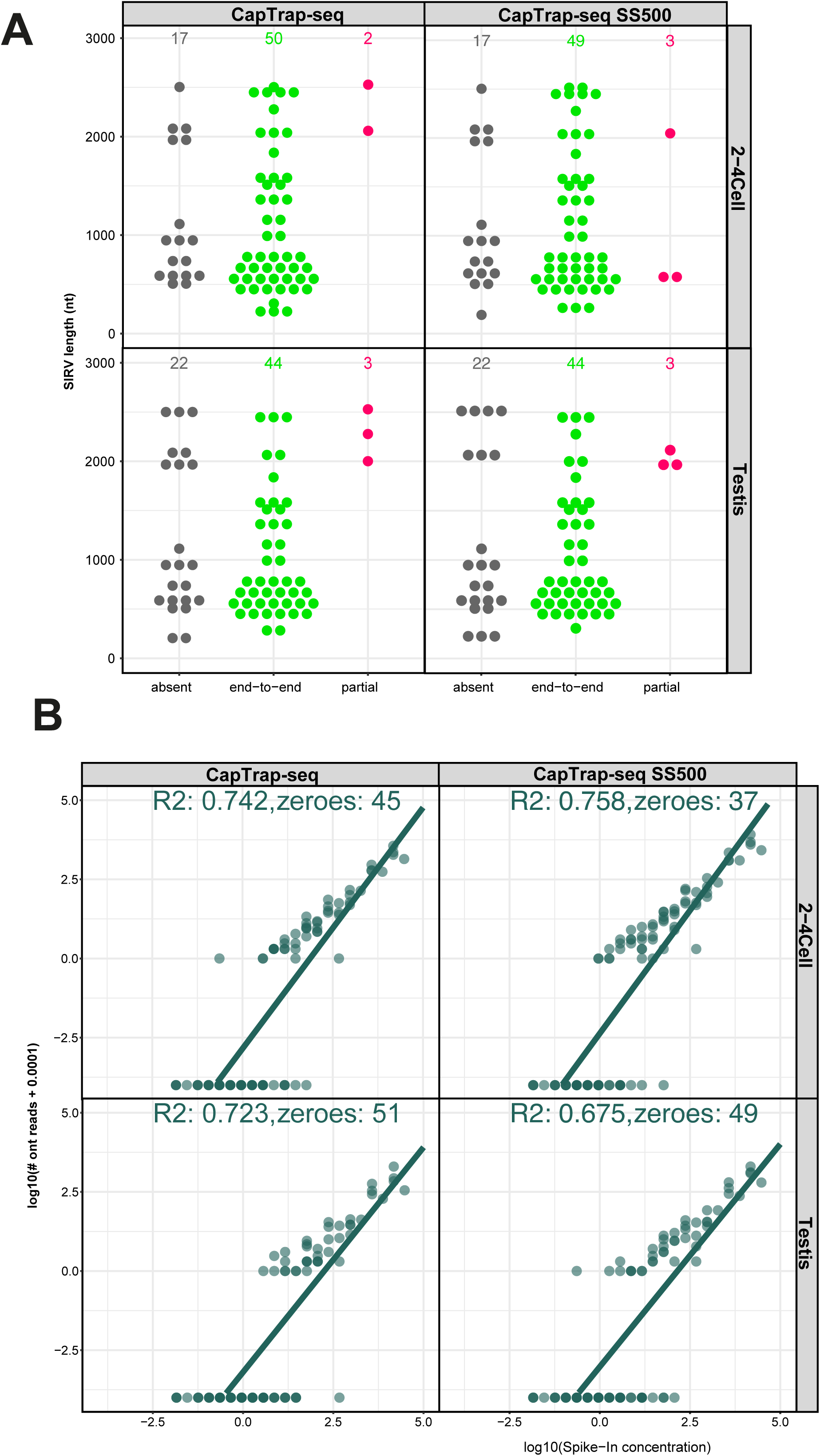
External RNA Spike-in detection and quantification analysis. **(A)** Detection of SIRVs in standard and size-selected (SS500) CapTrap-seq protocols as a function of transcript length. SIRVs are classified into three detection levels: end-to-end (green), partial (red), and undetected or absent (gray); **(B)** Relationship between input RNA concentration and raw read counts for ERCC spike-ins in 2-4 cell and testis samples. Each point represents an individual synthetic ERCC control. The green line represents a linear regression fit to the data, with the R^2^ value displayed at the top.

**Figure S10.**
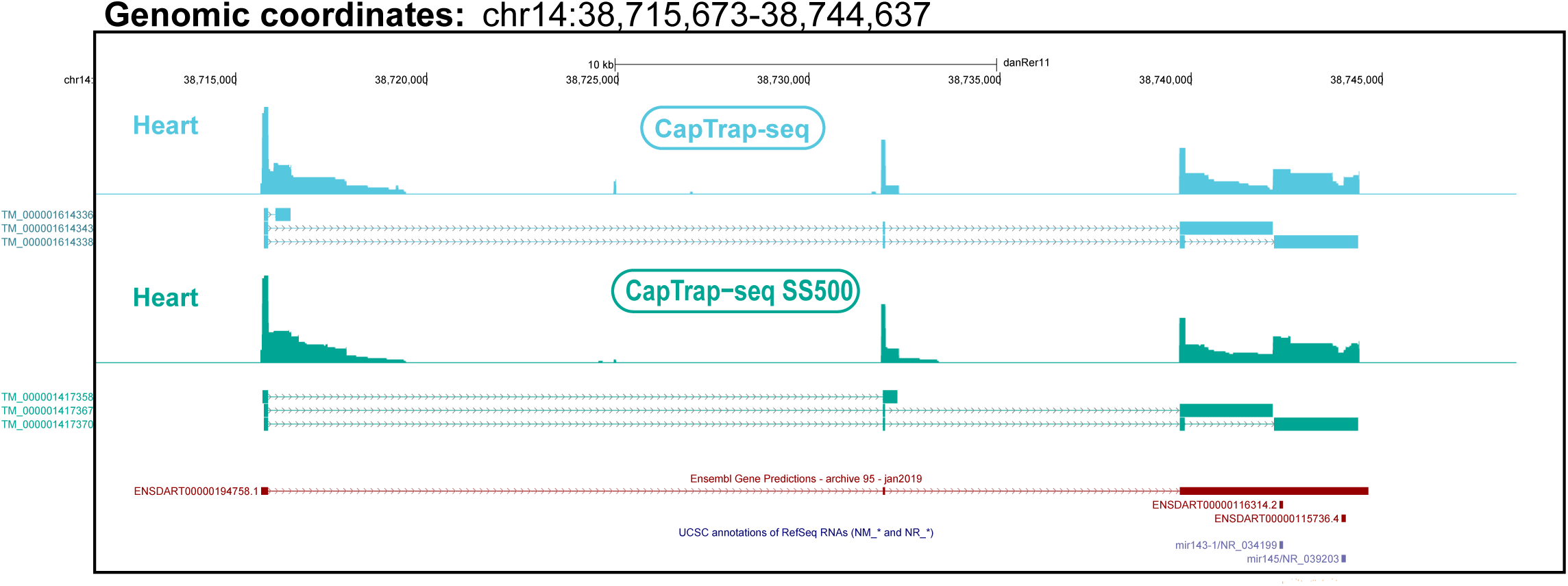
The effect of CapTrap-seq size selection on the identification of novel lncRNA transcripts. Novel transcript models identified for the *carmn* lncRNA gene in zebrafish. Colors represent the library preparation method: blue for CapTrap-seq and green for size-selected (SS500) samples . Ensembl models are displayed in burgundy, while RefSeq models are shown in navy blue. The corresponding bigwig files from long-read ONT RNA-seq data are shown above each transcript model, utilizing signal track mode in full view on the UCSC Genome Browser. Vertical black lines mark 5′ end support from DANIO-CODE^31^ CAGE-seq signals and 3′ end support from DANIO-CODE 3P-seq data.

